# Gene Expression Profiling and Physiological Adaptations of Pearl Spot (*Etroplus suratensis*) under Varying Salinity Conditions

**DOI:** 10.1101/2023.05.24.542058

**Authors:** Pranali Prabhakar Marbade, S. A. Shanmugam, E. Suresh, A. Rathipriya, Deepak Agarwal

## Abstract

*Eutroplus suratensis* (Pearl spot) is naturally found in estuarine environments and has been noted to have a high salinity tolerance, with the ability to thrive in freshwater as well as seawater. By examining the impact of various salinity levels on the growth and survival of Pearl spot, the present study aims to enhance aquaculture profitability by assessing their adaptability and physiological adjustments to changes in salinity, as well as determining their potential to acclimate to a broad range of salinity regimes. Pearl spot fingerlings were placed in tanks with varying salinities (15, 25, 35, 45, 60, and 75ppt) and monitored for mortality at 24-hour intervals up to 120 hours. Results revealed no mortality in the control group (0ppt), as well as in the 15, 25, and 35ppt treatment groups. However, the remaining groups (45, 60, and 75ppt) showed differing levels of mortality, with 44% mortality observed in the 45ppt group and 100% mortality in both the 60 and 75ppt groups. The impact of different salinity levels on the expression of pearl spot genes such as IGF-1, SOD, CAT, NaKATPase, OSTF-1, and HSP70 was investigated, along with a histological examination of the gills. The results showed significant physiological and cellular damage caused by the salinity levels. The expression analysis showed that liver IGF-1 mRNA expression increased by 2.6-fold at 15ppt, and HSP70 mRNA expression in the liver also showed a significant increase with rising salinity levels. In addition, OSTF1 expression exhibited an increase at 15ppt, whereas SOD and CAT expression reached their highest levels at 25ppt. At 15ppt, the expression of NKA mRNA increased significantly by 2.8-fold. The study’s overall findings suggested that the fish demonstrated strong molecular-level performance between 15 to 25ppt salinity levels, with the best results observed at 15ppt. These findings suggest that utilizing a salinity level of 15ppt for Pearl spot production could be viable for profitable aquaculture.

## 1. Introduction

The impact of climate change on aquatic organisms has become an area of increasing concern due to the potential implications for biodiversity and ecosystem functioning. Climate change is exerting varying degrees of physiological stress on the fishes by knocking the metabolic pathways closely associated with homeostasis and physiological adaptation to the changing environment, especially temperature and salinity fluctuations. Salinity is an important physical factor that affects many aspects of physiology such as growth, reproduction, osmoregulation, etc. and has pervasive effects on physiological and biochemical functions at all levels of biological organization, from molecule to organism. According to the tolerance range of salinity, the fishes can be classified into two groups, euryhaline and stenohaline species. The pearl spot fish (*Etroplus suratensis*) is an economically important species in South Asia that inhabits estuarine and freshwater environments. This species can tolerate a wide range of salinities by using efficient osmoregulation to maintain homeostasis (Evans et al. 2005) and highly tolerated cellular stress response (Chandrasekar *et al.,* 2014), making it an ideal model for studying the effects of salinity on gene expression and growth patterns. Salinity exposure is known to affect growth performance (Kang’ombe and Brown, 2008; Ninh et al., 2015; Gan et al., 2016), reproductive capacity (Vieira et al., 2019), digestive capacity (Tran-Ngoc et al., 2017), blood parameters (Verdegem et al., 1997), immune function parameters (Choi et al., 2013), antioxidant status (Gan et al., 2016), plasma osmolality, respirometry response (Morgan et al., 1997), metabolicrate (Iwama et al., 1997), histopathology and behaviour (Hassan et al., 2013). Despite the negative effects of salinity, some reports witnessed growth improvement in seawater-reared freshwater fishes under suitable salinity ranges (Sparks et al., 2003). The world has been experiencing the effects of global warming as a result of the accumulation of carbon dioxide in the environment. The unfavourable impacts of global warming enhance the temperature of the earth’s atmosphere, significantly influencing salinity (Fernández et al., 2022). Aquatic life is impacted by shifts in salinity in terms of physiological behaviour, adaptability, growth, and breeding pattern. Along with global warming, several changes to the saltwater ecology, such as the construction of a barrage for rice farming, the rapid extension of backwater tourism, the removal of mangroves, etc., are also responsible for the rapid decline of the pearl spot in native habitat (Padmakumar. et. al 2012). Many aquatic organisms have found it difficult to adapt and survive in these conditions due to the wide range of salinity fluctuations that alter the physiological conditions in the coastal and estuaries.

Although the pearl spot fish has been studied at various salinities (Chandrasekar et al., 2014), the fish has not yet been examined at the molecular level from the perspectives of adaption and growth at various salinities. This study aims to identify and characterize changes in gene expression within the major osmoregulatory tissues of the pearl spot. Recent research has focused on the analysis of gene expression in pearl spot fish adapted to different salinities to better understand the mechanisms underlying their adaptation. By studying the expression of genes related to growth, osmoregulation, ion transport, and other physiological processes, researchers hope to gain insight into how pearl spot fish can survive and thrive in changing salinity environments. The kidney, intestine, and gills are the main group of osmoregulatory organs for regulating ion movements between gain and loss. In the gill epithelium, there are mitochondria cells (MR), which are in charge of the uptake of the ions in freshwater fish (FW) and the ions secretion in saltwater fish (SW) (Hiroses et al., 2003). To maintain homeostasis, the epithelial cells in gills can withstand a variety of salinities and osmotic stresses (Arun Kumar et al., 2020). Histological analysis of gills can provide insights into the structural changes that occur during adaptation to different salinity levels. Therefore, in this study, we also examined the histological changes in the gills of pearl spot fish during adaptation to various salinity levels, intending to gain a better understanding of the physiological mechanisms involved in salinity adaptation in this species. Understanding the molecular and cellular mechanisms underlying salinity adaptation in pearl spot fish can inform the development of strategies for managing fish populations in response to climate change. Aquaculture may be more profitable if species can adapt to a variety of salinity levels.

## 2. Materials and Methods

### 2.1. Sample and Tissue Collection

The wet lab experiments were conducted at TNJFU – Institute of Fisheries Post Graduate Studies, OMR Campus, Vaniyanchavadi, Chennai. Molecular analysis was performed in the Molecular Biology Laboratory of TNJFU-IFPGS, Chennai. The experimental fishes (fingerlings of pearl spot of around 5-7 cm) were procured from ICAR-CIBA, Muttukadu centre, Chennai and acclimatized in wet lab conditions for 2 weeks in normal bore well water at IFPGS.

To determine the LC50 for salinity, groups of nine fishes were reared with triplicate at varying salinity levels including a control (0ppt), and their mortality was observed every 24 hours for 120 hours. For the adaptation study, groups of twenty fishes with triplicate were reared at 0ppt for 7 days, after which sampling was done on the 8th day. The salinity was then gradually increased by 1-2ppt per day until reaching 15ppt, at which point the fishes were reared for an additional 7 days before sampling on the 8th day. This process was repeated for salinity levels of 25, 35, and 45ppt, with gradual increases in salinity and 7-day rearing periods followed by sampling on the 8th day. Gill, kidney, and liver tissues were collected from each fish in each replicate at each fixed salinity, and all the sampled tissues were stored in RNAlater^TM^ (QIAGEN, USA) at -80°C for subsequent analysis.

### 2.2. Histological Analysis

To investigate the impact of salinity on a cellular level, histology was conducted on the gills of the pearl spot. The fish tissues were removed and immersed in 10% neutral buffered formalin, then stored in the same buffer at room temperature. Subsequently, the tissues were dehydrated with increasing concentrations of ethanol, cleared with xylene, and embedded in paraffin wax with a congealing point of 58-60°C. Thin sections of 6-8 μm were obtained from the paraffin block using a rotary microtome from Leica, Germany, and mounted on albumenised slides. The slides were fixed overnight at 60°C, and the sections were deparaffinized with xylene and dehydrated with decreasing concentrations of alcohol to distilled water. After staining with haematoxylin for 20 minutes, the sections were differentiated in 1% alcohol and treated with ammonia water, washed, and then stained with 2.5% w/v eosin for 10 minutes. The sections were then dehydrated, cleared, and mounted in DPX (Distrene, Plasticiser, Xylene), and physiological changes under varying salinity conditions were observed using an Olympus microscope.

### 2.3. Total RNA Isolation and Reverse Transcriptase PCR (RT - PCR)

Total RNA was extracted from Gill, Liver and Kidney of Pearl spot sing TRIzol^TM^ reagent (Invitrogen, USA) following Sambrook et al. (2001). The integrity of the isolated RNA was checked using the electrophoresis method on 1% agarose gel. The purity and quantity of the extracted RNA were verified with a Nanodrop spectrophotometer (Thermoscientific, USA), which directly provided the total concentration in terms of ng/μL along with the 260:280 ratios. The ratio of the absorbance at 260 nm and 280 nm provided the estimate of the purity of the isolated RNA. To eliminate genomic DNA contamination, RNase-free DNAse I (Thermoscientific, USA) was utilized to treat the extracted total RNA. The RNA samples were then stored at -80°C for further use. The mRNA from all the extracted RNA was converted to its complementary DNA using the RevertAid First Strand cDNA Synthesis Kit (Thermoscientific, USA) as per the manufacturer’s instructions. The cDNA was diluted at a 1:10 ratio and was used for the real-time PCR (qRT-PCR).

### 2.4. Primers Designing and PCR Amplification

Except for SOD and Catalase, the nucleotide sequences of all targeted genes of pearl spot were obtained from NCBI. The Gene-specific primers (Table 1) for all the selected genes were designed using Gene Runner software (version 6.5) and Primer3 Plus (Untergasser *et al.,* 2007). For SOD and Catalase, the nucleotide sequences were not reported at NCBI for pearl spot, hence degenerated primers were designed from the consensus sequences of closely related cichlids such as Tilapia, sea bass, etc. Several factors, such as primer length, % GC content, melting temperature, annealing temperature, secondary structures, etc. were taken into consideration while designing the primer sets. A total of 7 primer pairs were designed and commercially synthesized by Gene to Protein (Biotech desk), Chennai. The annealing conditions for all primer combinations were optimized through gradient PCR (Table 1). For PCR, 25μl reactions were set up, including 12.5μl Taq DNA Polymerase Master Mix RED (Ampliqon, Denmark), 1μl of each primer (10 pm) and 1μl cDNA (70ng/μl). The program comprised initial denaturation at 94°C for 3 minutes, followed by 35 cycles of denaturation at 94°C for 45 seconds, annealing at the appropriate temperature for 45 seconds, and extension at 72°C for 45 seconds. A final extension step was performed at 72°C for 5 minutes.

**Table 1.**
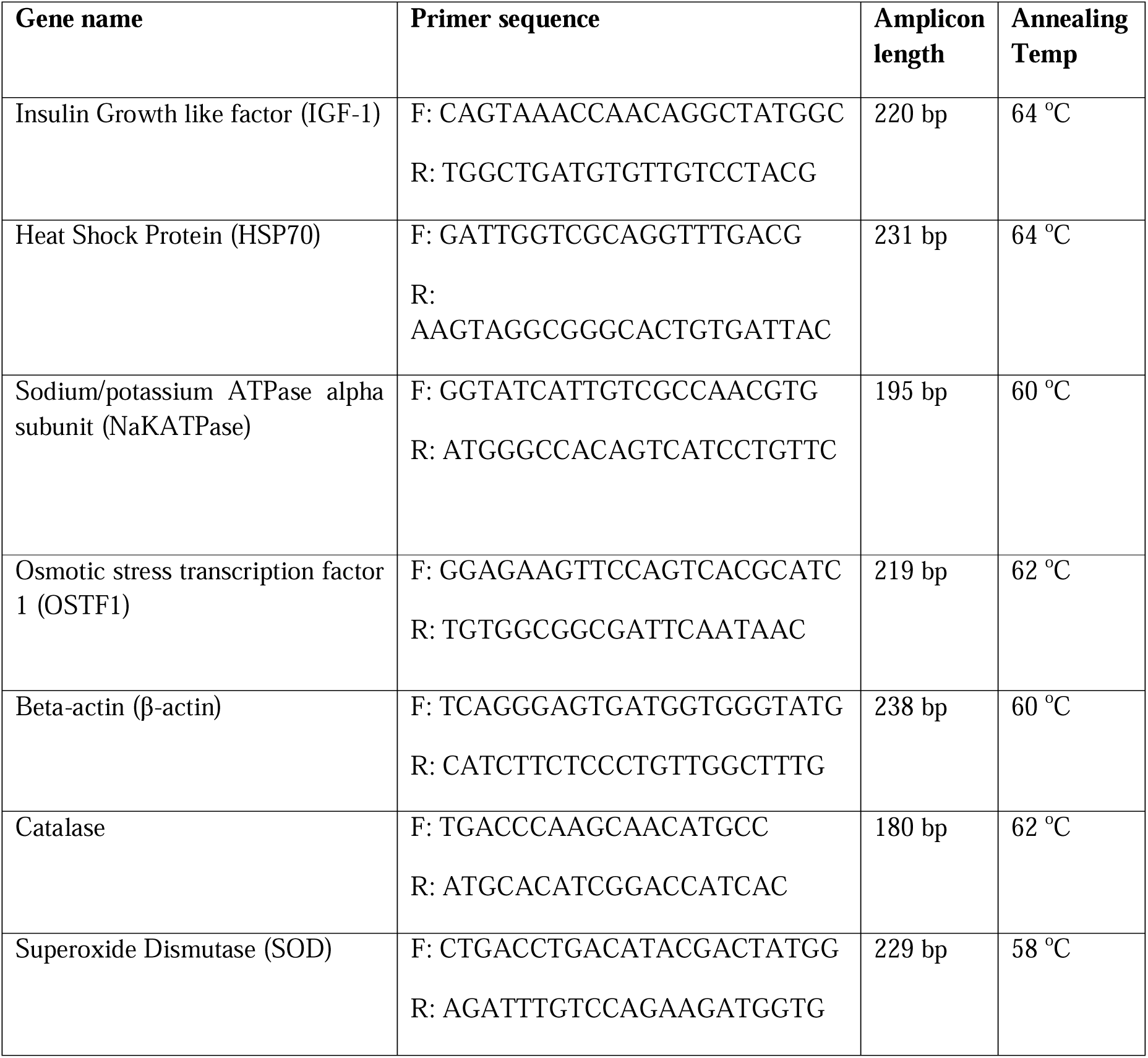
Primer and PCR amplification details of selected genes.

### 2.5. Gene Expression Analysis by Real-time PCR

Quantitative real-time PCR was conducted using SYBR Green chemistry on a Quant Studio-5 instrument (Applied Biosystems, USA). The reaction volume of 20 µL consisted of 10 µL of TB Green Premix Ex Taq II (Tli RNase H Plus) (Takara, Japan), 0.8 µL each of gene-specific primers (10 µM), 0.4 µL of ROX Reference Dye (50X), 2 µL of cDNA (20 ng), and 6 µL of nuclease-free water. The PCR cycling conditions included an initial denaturation step at 95°C for 30 sec, followed by 40 amplification cycles of denaturation at 95°C for 5 sec and annealing at gene-specific temperature for 34 sec, and a melt curve step of 95°C for 15 sec, 60°C for 1 min, with a continuous temperature increase until 95°C and a 15 s hold at dissociation. All reactions were performed in triplicate and repeated thrice. The relative expression of genes was analyzed using the comparative Ct method (2^−ΔCt^) (Livak et al., 2001). Two putative housekeeping genes (HKG), namely Elongation Factor 1-alpha (EF1α) and β-actin, were statistically analyzed using the comprehensive algorithms of NormFinder, BestKeeper, and Genorm to assess their transcriptional expression stability. After analysis, β-actin was selected as the HKG for gene normalization, considering its stability and suitability for the analysis.

### 2.6. Statistical Analysis

The data were subjected to a one-way analysis of variance (ANOVA) followed by Duncan’s multiple range tests using the statistical package SPSS 29.0 computer program (SPSS Inc., Chicago, IL). The value P < 0.05 was considered statistically significant. All values were expressed as the means ± standard error of the mean.

## 3. Results

### 3.1. Histological Analysis

To investigate the impact of osmotic stress on fish at the cellular level, the histological analysis of *Etroplus surantensis* gills was conducted at five different salinity levels: control (0ppt), 15, 25, 35, and 45ppt. The gill structure of the control fish appeared normal, with chloride cells primarily located at the base of the interlamellar space (Fig 1). At 15ppt salinity, there were no significant histological changes observed in the fish gills, but those grown at 25ppt displayed certain physiological alterations. High salinity levels affected the density and size of chloride cells, as well as induced edema and hypertrophy in the lamellar region of the gill, leading to congested blood vessels (Fig 2 & 3). Overall, these changes indicated the occurrence of osmotic stress at 25ppt. At 35 and 45ppt, several cellular damages were observed such as lamellar curling, hyperplasia, and ruptured epithelia with haemorrhages. In addition, chloride cells in the lamellar regions grew larger due to lamellar fusion and necrosis induced by extreme salinity. (Fig 4 & 5).

**Fig. 1.**
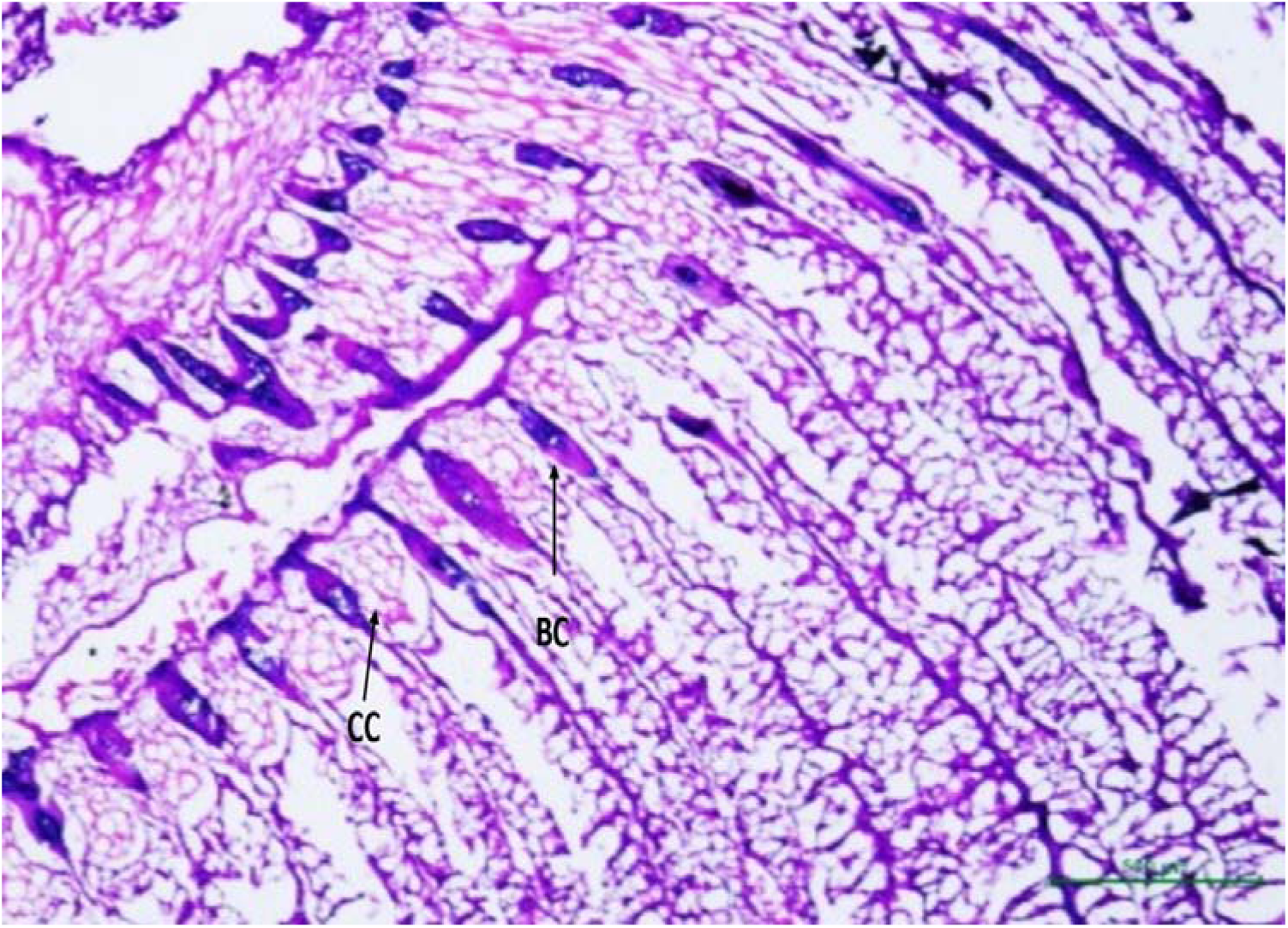
Histological section of gill tissue from *Etroplus suratensis* fish reared at 0ppt salinity. The figure shows the normal gill structure of the control fish. Chloride cells are predominantly found towards the interlamellar space’s base. CC: Chloride cells, BC: Blood Channels (Scale bar: 500μm, objective 4X).

**Fig. 2.**
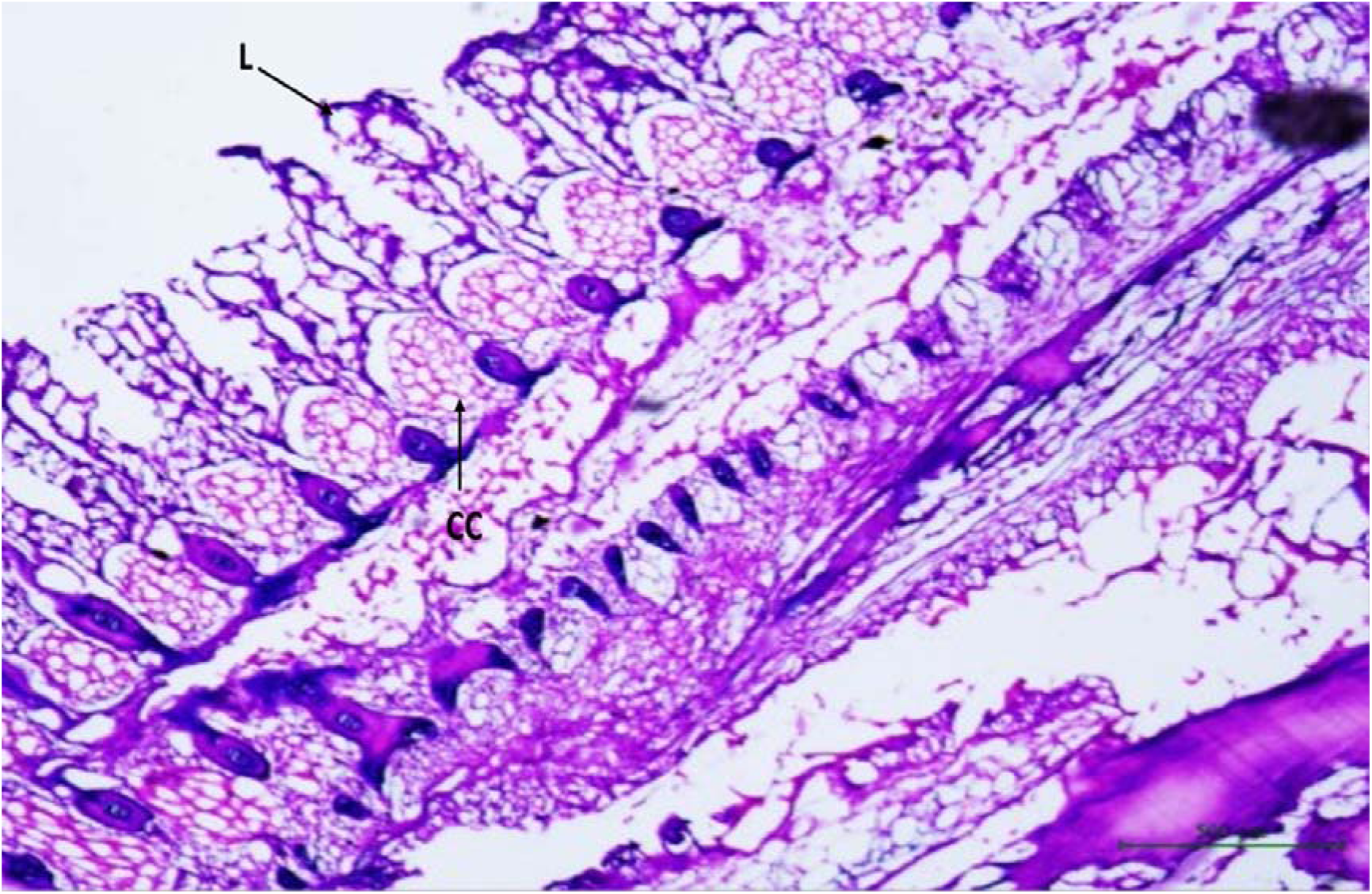
Histological section of gill tissue from *Etroplus suratensis* fish reared at 15ppt salinity. No significant histological changes were observed compared to the control group. CC: Chloride cells, L: Lamellar (Scale bar: 500μm, objective 4X).

**Fig. 3.**
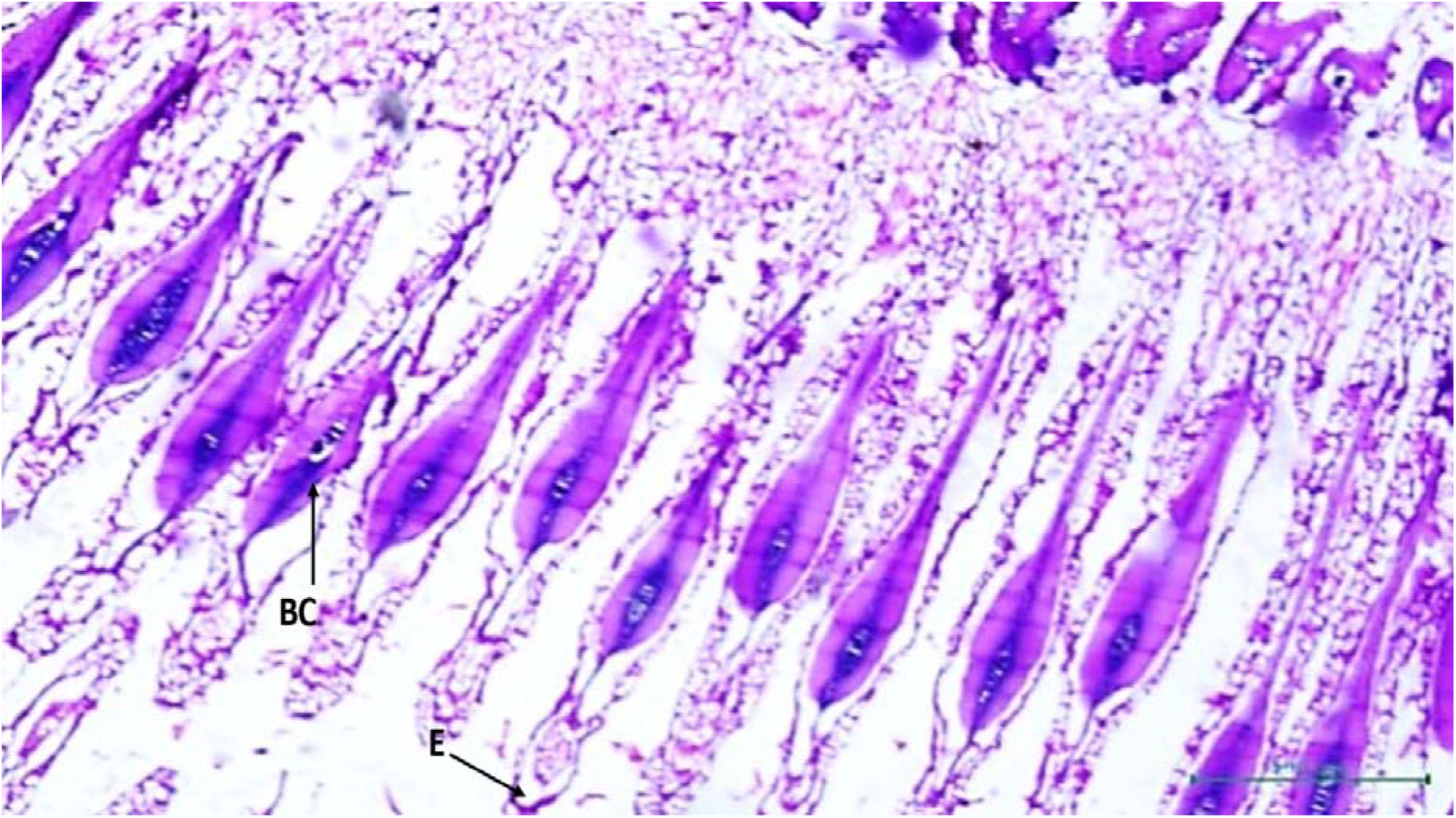
Histological sections of *Etroplus surantensis* gills at 25ppt salinity showing physiological changes. The chloride cells’ density and size were affected, and the lamellar region of the gill was similarly affected by edema and hypertrophy due to salt. Blood vessels were congested, and there were physiological adaptations due to osmotic stress. E: Edema (Scale bar: 500μm, objective 4X).

**Fig. 4.**
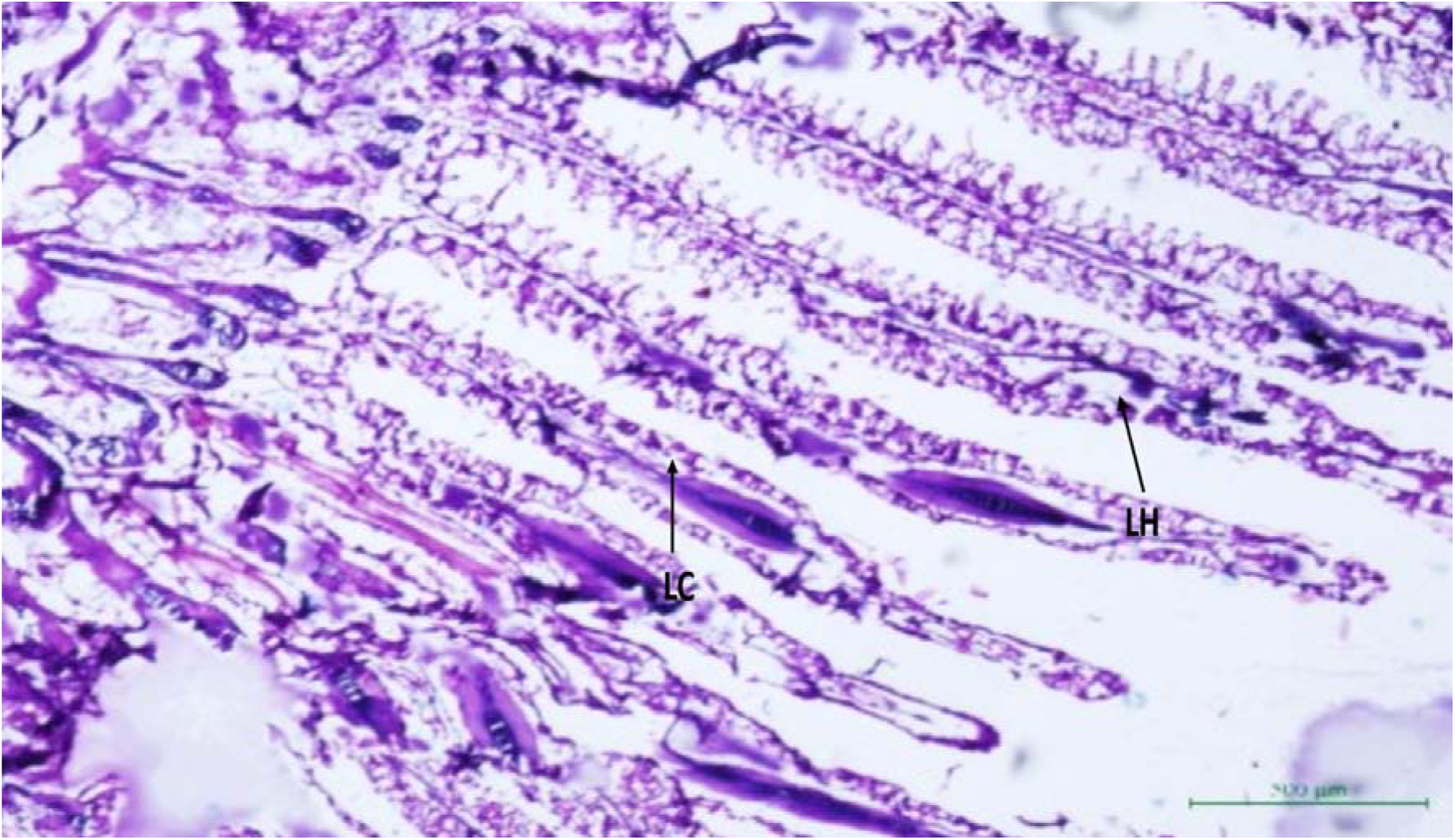
Histological section of *Etroplus suratensis* gills at 35ppt salinity. Lamellar curling, hyperplasia, and rupture epithelia with haemorrhages are evident. LC: Lamellar Curling, LH: Lamellar Hyperplasia (Scale bar: 500μm, objective 4X).

**Fig. 5.**
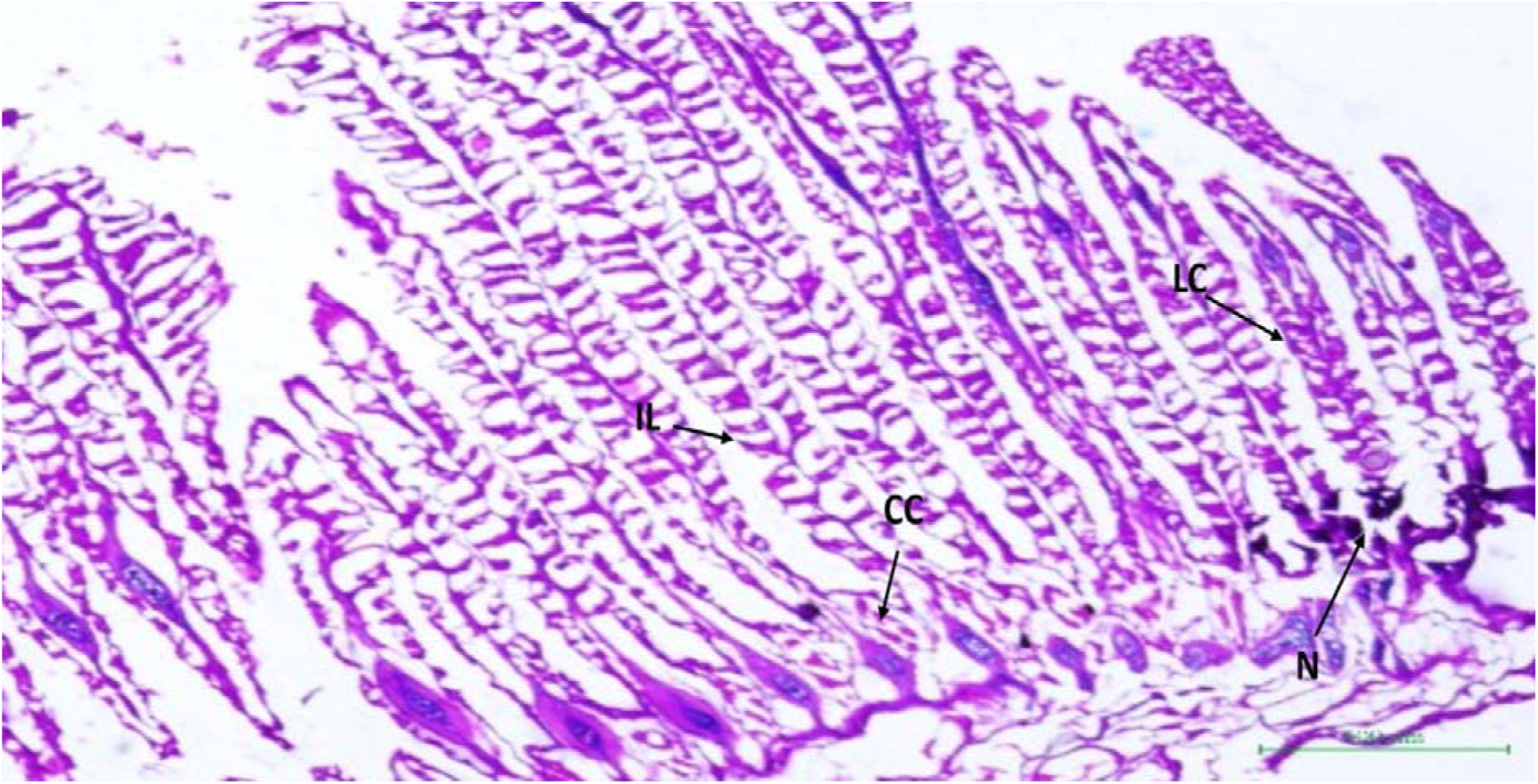
Histological section of *Etroplus suratensis* gills at 45ppt salinity. The chloride cells are notably enlarged and found in lamellar regions due to lamellar fusion and necrosis induced by extreme salinity. N: Necrosis, IL: Inter-lamellar, LC: Lamellar Curling. (Scale bar: 500μm, objective 4X).

### 3.2. LC50 for salinity in Eutroplus suratensis

The adaptability and potentiality of a species to different salinity regimes were investigated through an LC50 study. Fish were exposed to salinities ranging from 15 to 75ppt including a control (0ppt) to determine the upper limit of salinity exposure. Mortality was monitored at 24h intervals over 120 hours for each salinity treatment, and LC50 values were calculated using the probit method in SPSS software. No mortality was observed in the control and salinity treatments of 15, 25, and 35ppt, while hypersaline treatments (45, 60, 75ppt) exhibited varying levels of mortality. A 44% mortality rate was observed at 45ppt salinity exposure, whereas 100% mortality was recorded at 60 and 75ppt salinity treatments. The LC50 value for salinity was determined to be 45.394 ppt (Fig 6).

**Fig. 6.**
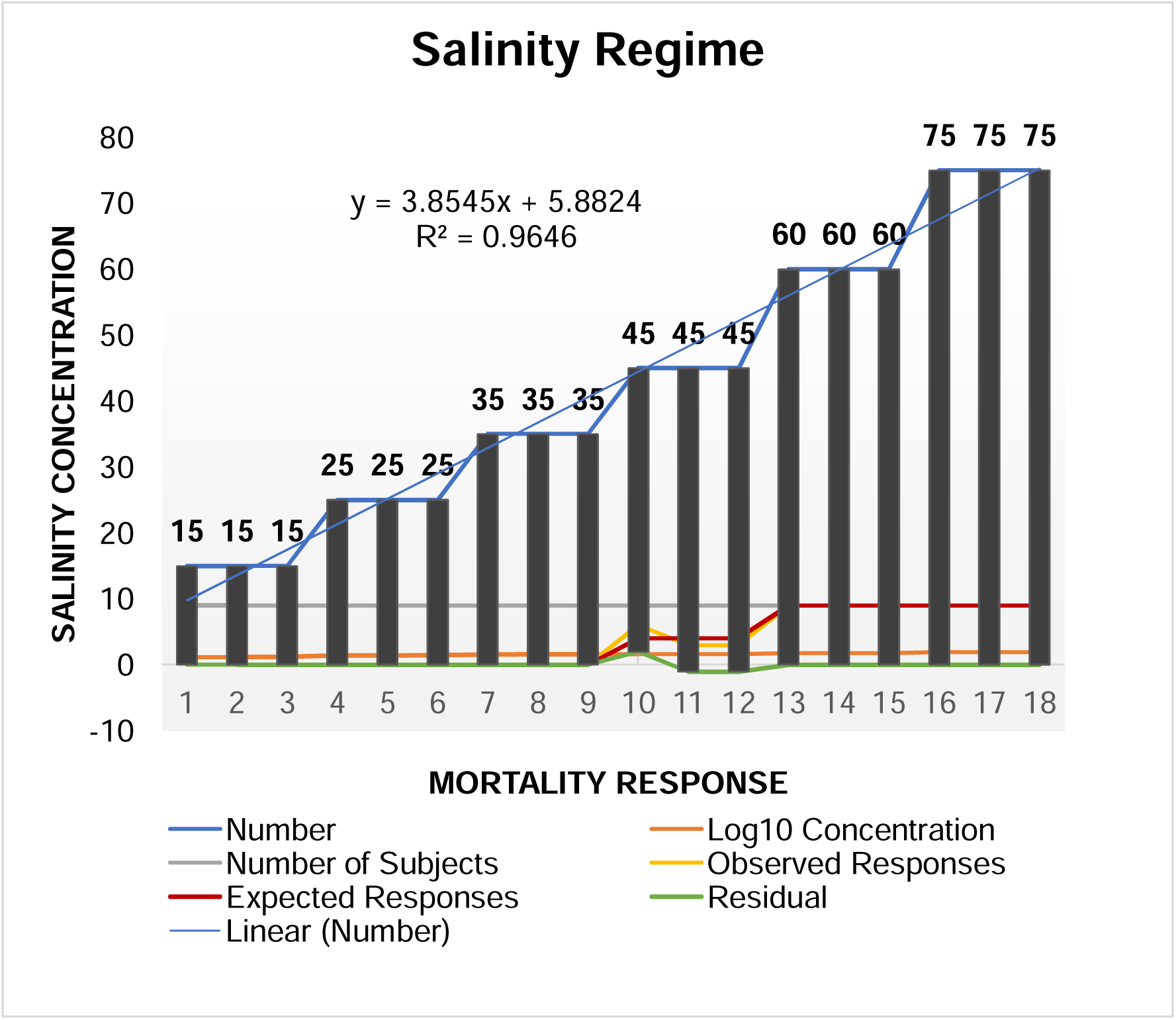
Probit graph showing the relationship between salinity exposure and mortality in *Etroplus suratensis*.

### 3.3. Gene Expression Analysis using Real-Time PCR

To determine the fold change of a target gene, its CT value was compared to that of the endogenous control gene β-actin, which served as the internal control. CT values were analyzed, and ΔΔCt values were calculated using the internal and endogenous controls. Fold change was then calculated using the formula 2^(–ΔΔCt), and all resulting values were subjected to statistical analysis using SPSS.

#### 3.3.1. Growth gene expression (Insulin-like growth factor - 1 (IGF-1))

The mRNA expression level of Insulin-like Growth Factor-1 (IGF-1) in the liver was analyzed under different salinity conditions. The results showed a significant increase in IGF-1 mRNA expression at 15ppt salinity, with a fold change of 2.6 compared to the control group. On the other hand, a significant decrease in IGF-1 mRNA expression was observed as the salinity increased. At 45ppt, the mRNA expression of IGF-1 was significantly downregulated, with a fold change value of -6.4 compared to the control (as illustrated in Fig. 7). These observations indicate that salinity has a significant impact on the regulation of IGF-1 mRNA expression in the liver.

**Fig. 7.**
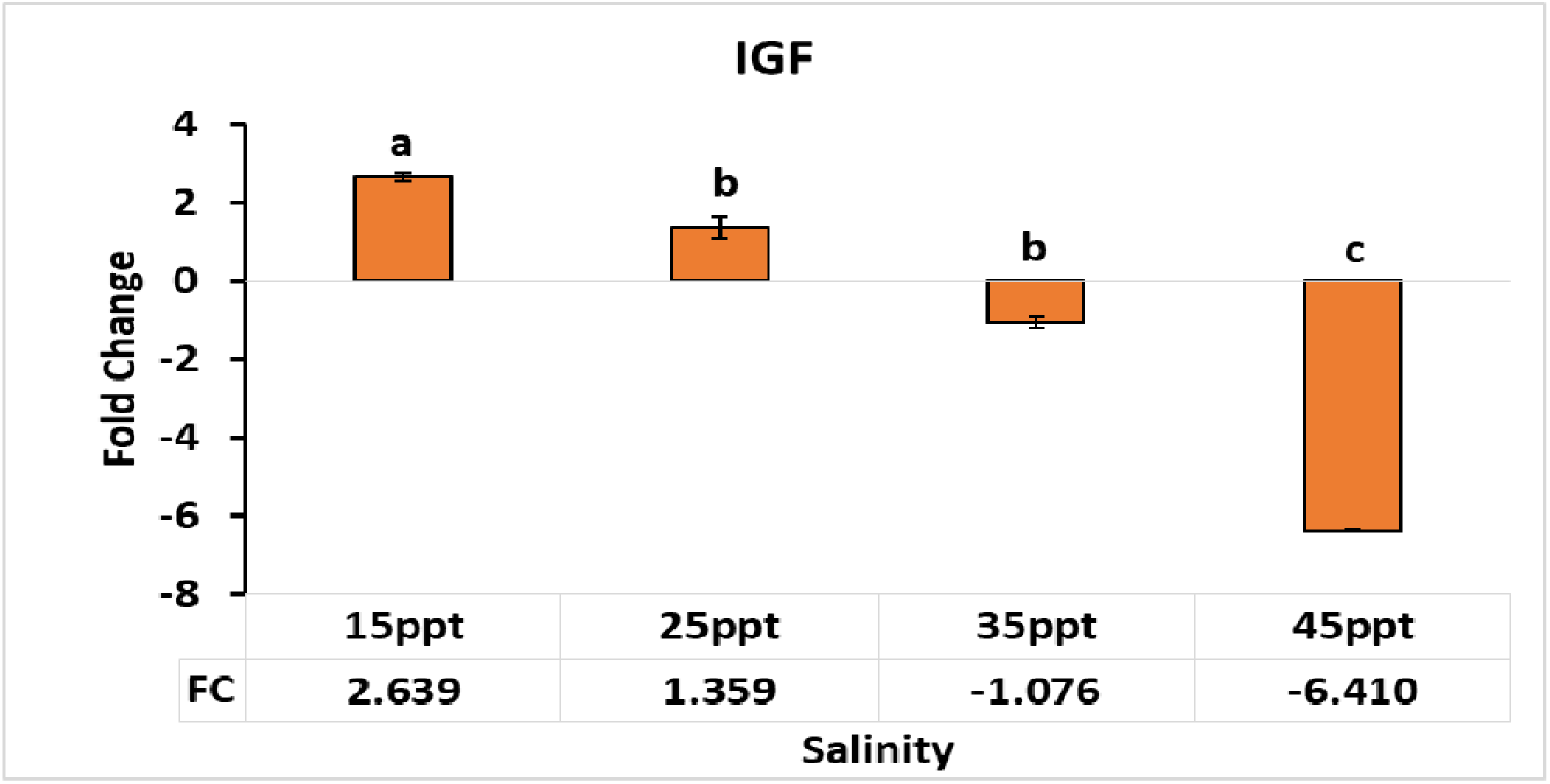
Fold change of IGF-1 mRNA expression in the Liver of *Etroplus suratensis*.

#### 3.3.2. Oxidative stress related gene expression (Heat shock protein 70(HSP70))

The HSP70 mRNA expression level was investigated in the liver of pearl spot at different salinities. The results showed a significant increase in HSP70 mRNA expression levels at 15ppt salinity, compared to the control group (0ppt). Further increases in salinity levels were found to induce a progressive increase in HSP70 mRNA expression. Notably, the fold change values at 25 and 35ppt salinity were found to be statistically significant compared to the expression level at 15ppt salinity. The highest level of HSP70 mRNA expression was observed at the extreme hypersaline condition of 45ppt, with a fold change value of 7.3, as illustrated in Fig. 8. These findings suggest that HSP70 plays a crucial role in the adaptive response of pearl spot to salinity stress, and may have potential applications in aquaculture.

**Fig. 8.**
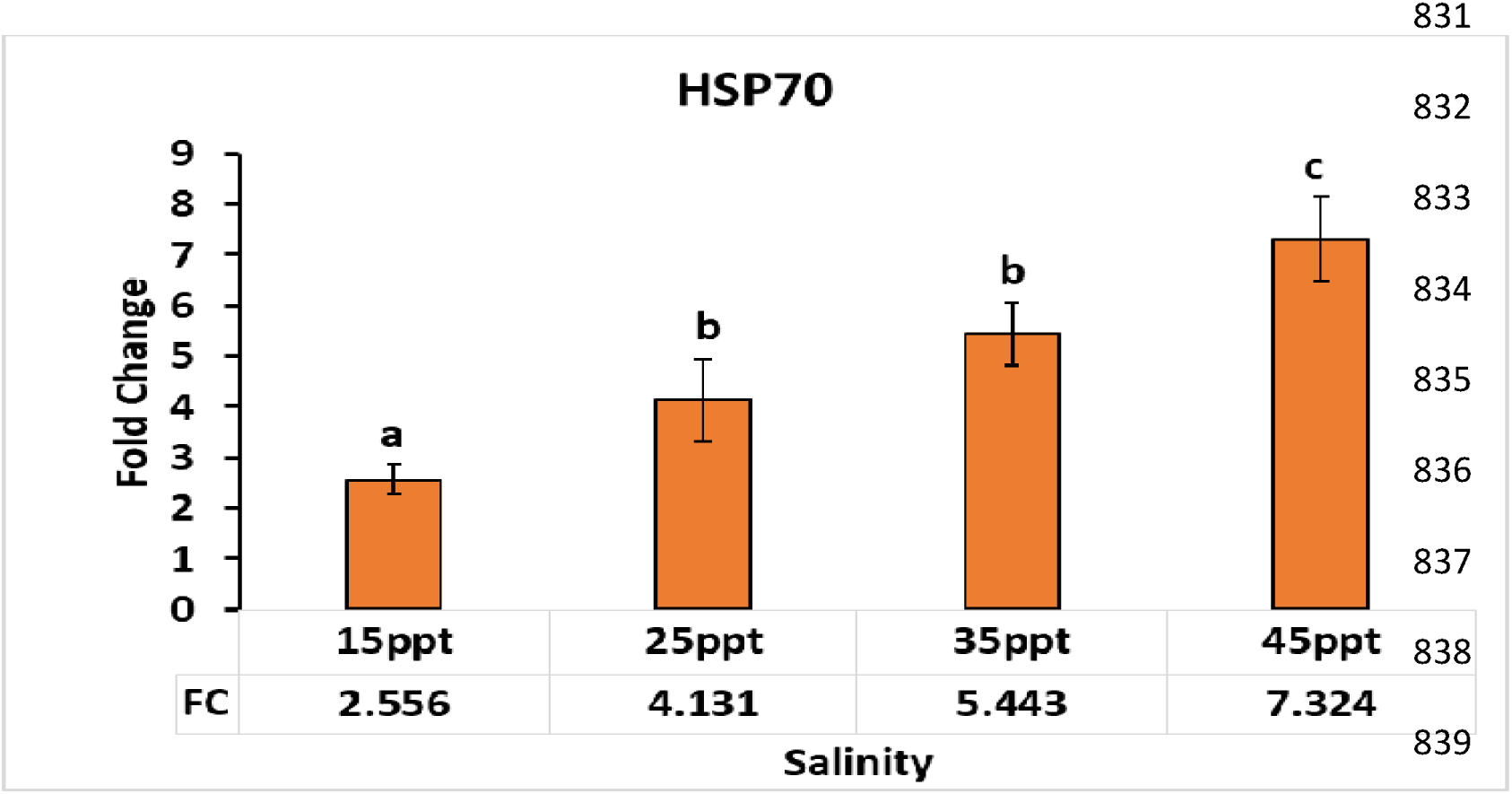
Fold change of HSP70 mRNA expression in the Liver of *Etroplus suratensis*.

#### 3.3.3. Osmotic stress related gene expression (osmotic stress transcription factor-1 (OSTF1))

The mRNA expression of osmotic stress transcription factor -1 (OSTF1) was investigated in the liver of pearl spot under different salinity conditions. The results revealed a gradual increase in OSTF1 mRNA expression levels with increasing salinity. Specifically, the expression of OSTF1 mRNA was found to be significantly increased at a salinity of 25ppt, with a statistically significant fold change value of 11.3, compared to the control group. However, a further increase in salinity to 35ppt resulted in a downregulation of OSTF1 mRNA expression relative to the expression at 25ppt. Interestingly, a subsequent increase in salinity to 45ppt led to a significant increase in OSTF1 mRNA expression with a fold change value of 4.9, as illustrated in Fig. 9. These findings suggest that OSTF1 plays a crucial role in the adaptive response of pearl spot to salinity stress, and its expression is intricately regulated by the salinity conditions. Further studies are needed to elucidate the underlying molecular mechanisms and potential implications of OSTF1 in this process.

**Fig. 9.**
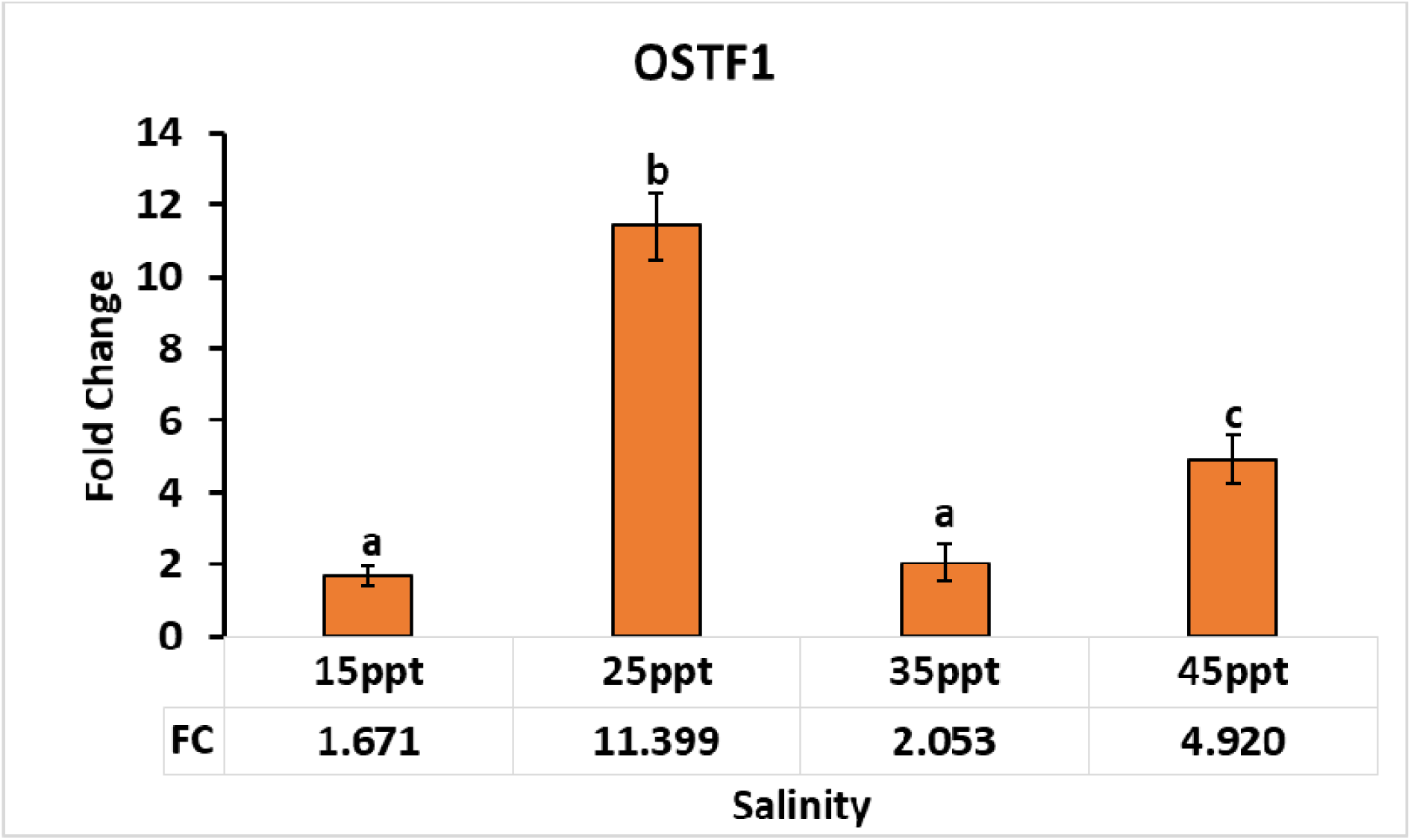
Fold change of OSTF1 mRNA expression in the Liver of *Etroplus suratensis*.

**Fig. 10.**
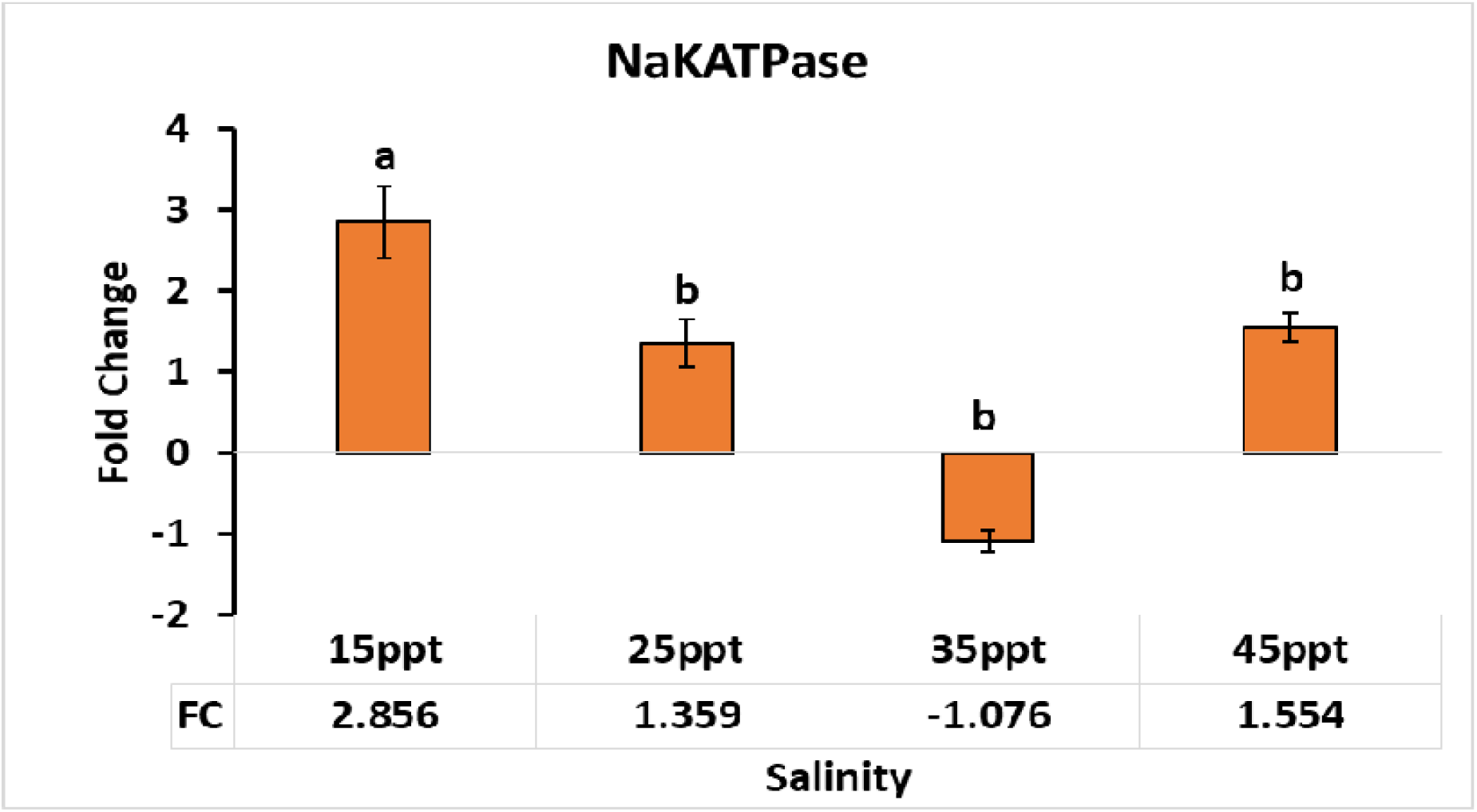
Fold change of NKA mRNA expression in the Gills of *Etroplus suratensis*.

#### 3.3.4. Ion regulatory gene expression (Sodium/potassium ATPase alpha subunit (NaKATPase))

The mRNA expression of Na+/K+-ATPase (NKA) was examined across different salinity levels in the gill tissue of the pearl spot. Findings showed that NKA mRNA expression was upregulated in response to increasing salinity levels. Notably, the highest level of NKA mRNA expression was observed at 15ppt salinity, with a statistically significant fold change of 2.8 compared to the control group. This finding is consistent with previous studies that have reported increased demand for ion transporters, such as NKA, in fish exposed to high salinity environments. However, contrary to our expectations, we found a significant decrease in NKA mRNA expression at 35ppt, with a fold change of -1.07, compared to the expression level at 15ppt, as illustrated in Fig. 9. This unexpected finding suggests that there may be a threshold level of salinity beyond which NKA expression is no longer required or may even be inhibited. Further investigations are needed to understand the underlying mechanisms involved in the regulation of NKA mRNA expression under different salinity conditions and the potential implications of these findings for fish physiology and aquaculture.

#### 3.3.5. Antioxidant Enzyme Superoxide dismutases (SODs) and Catalase Activity (CAT)

The present study investigated the mRNA expression of two antioxidant enzymes, SOD and CAT, in the liver tissue of pearl spot fish exposed to different salinity conditions. Results revealed a significant increase in SOD mRNA expression levels with increasing salinity, as compared to the control. The fold change value at 15ppt was 4.4, indicating a significant upregulation of SOD expression. The maximum fold change values were observed at 25 and 35ppt, with fold changes of 10.6 and 10.2, respectively, when compared to the control (Fig 11). These findings suggest that SOD plays a critical role in combating oxidative stress induced by salinity stress in pearl spot fish. Similarly, the mRNA expression of CAT was also investigated in the liver tissue of pearl spot fish. Interestingly, CAT mRNA expression was considerably upregulated at 25ppt, as compared to the control and other salinity regimes, with a remarkably high fold change value of 26.5 (Fig 12). These findings suggest that CAT may play a crucial role in mitigating oxidative damage induced by salinity stress in pearl spot fish. Further studies are warranted to explore the underlying molecular mechanisms of these antioxidant enzymes in response to salinity stress and their potential implications in the adaptation of pearl spot fish to varying salinity conditions.

**Fig. 11.**
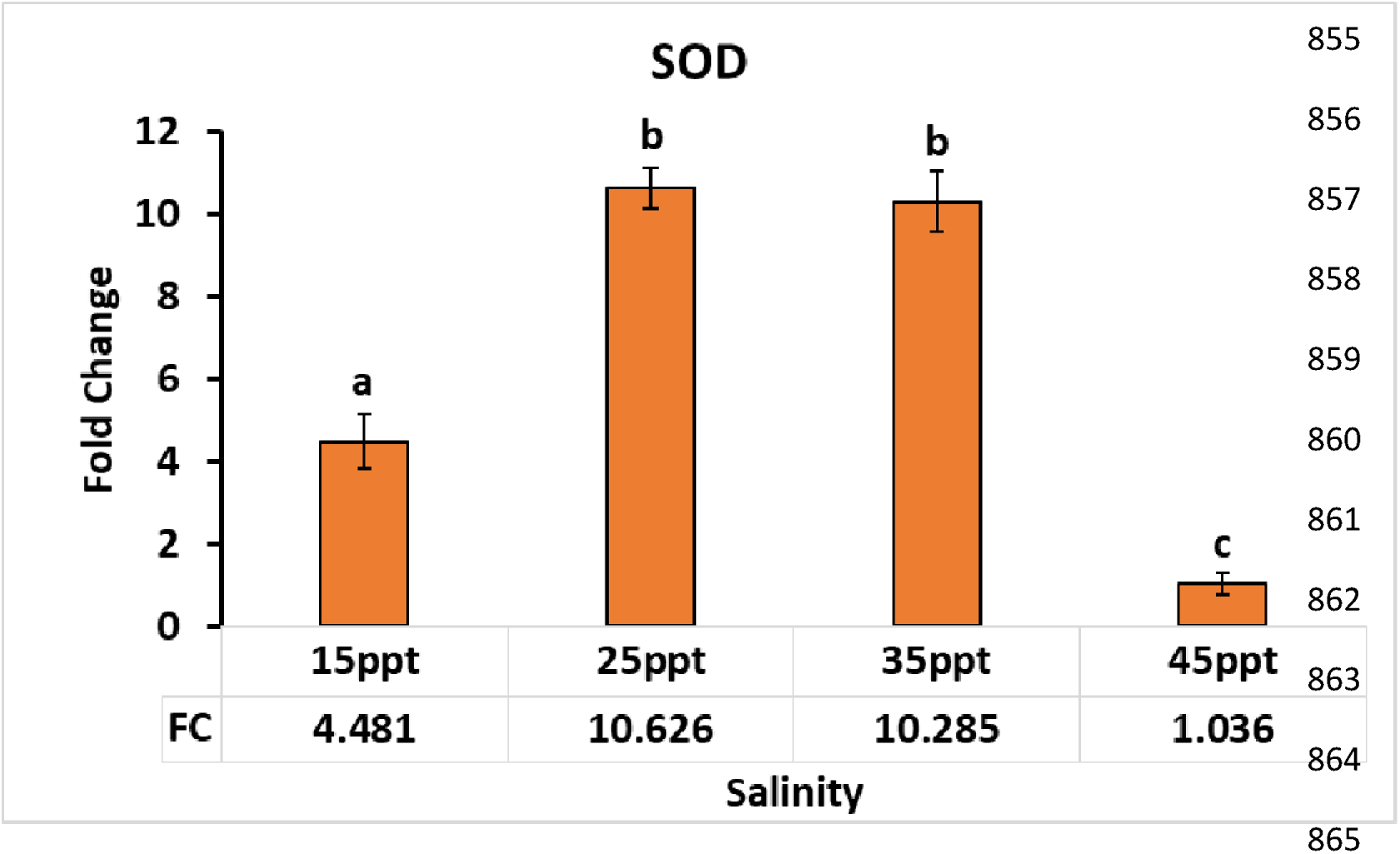
Fold change of SOD mRNA expression in the Liver of *Etroplus suratensis*.

**Fig. 12.**
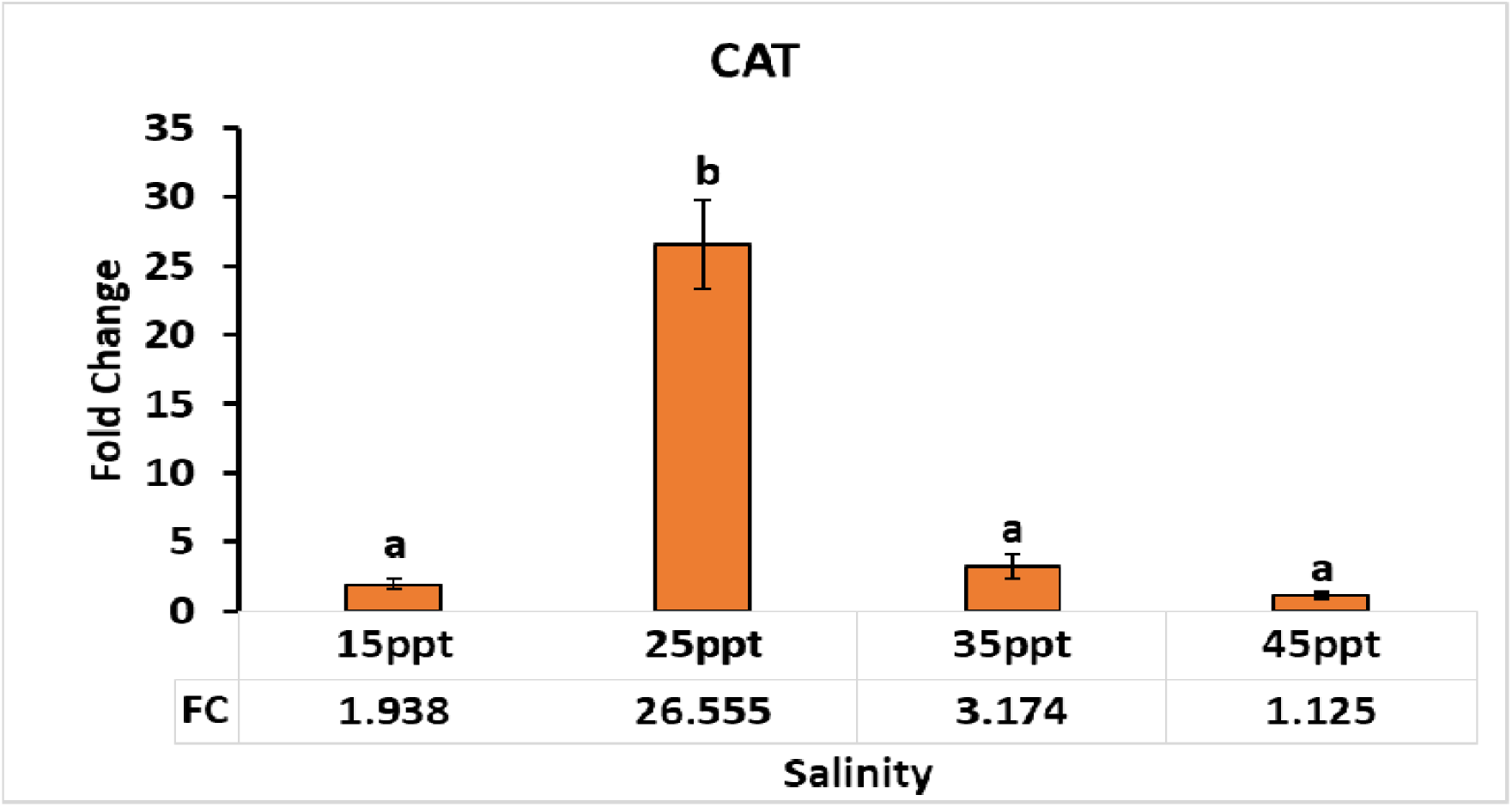
Fold change of CAT mRNA expression in the Liver of *Etroplus suratensis*.

## 4. Discussion

The impact of climate change-related salinity intrusion on coastal areas is a significant concern worldwide. To address this concern and understand the effects of varying concentrations of salinity on *Etroplus suratensis*, a species commonly known as pearl spot, the current study aimed to investigate the expression of critical genes related to different physiological processes, including growth, ion transportation, and stress response, at different salinity levels. In addition, to determine the upper limit of salinity tolerance for pearl spot, a lethal concentration study, commonly known as LC50, was conducted. This study aimed to identify the concentration of salinity that could cause at least 50% of the tested population to perish. Osmotic stress caused by high salinity levels can have detrimental effects on fish physiology, which can be further understood through histological studies. Therefore, it is crucial to investigate how different salinity levels impact the expression of these genes and to study the histological changes that occur in response to osmotic stress. Overall, this study highlights the critical importance of investigating the effects of salinity on fish species and their physiology, especially in the context of climate change-related salinity intrusion in coastal areas. By understanding the lethal concentration of salinity and the underlying physiological changes in response to osmotic stress, we can develop strategies to mitigate the impact of climate change on fish populations and ensure the sustainability of our oceans and coastal ecosystems.

In this study, *Etroplus suratensis* showed 100% survival at salinities of 0, 15, 25, and 35ppt, while exhibiting 100% mortality at 60 and 75ppt. At 45ppt, the fish showed a survival rate of only 44%. Based on the survival and mortality data, probit analysis determined the LC50 value of 45.394ppt for salinity in the pearl spot. Bringolf et al. (2005) conducted a study to determine the salinity tolerances of flathead catfish *Pylodictis olivaris*, a species with a broad habitat range, characterized by rapid dispersion within and among river systems, and the potential to migrate using seawater. The researchers reported that the LC50 values were 10ppt when the fish were exposed to NaCl and 14ppt when exposed to synthetic seawater. Another widely distributed fish, *Cichlasoma urophthalmus*, showed an LC50 value of 24ppt, with 100% survival occurring at 10ppt (Martinez-Palacios et al., 1990). In a study by Martinez et al. (1996), the LC50 value for salinity in the stenohaline freshwater Central American cichlid (*Cichlasoma synspilum*) was recorded as 14.5ppt after 144 hours. Akther et al. (2009) conducted a salinity tolerance trial on silver barb and reported an LC50 value of 14.19ppt at 96 hours, using common salt as the salinity source. Sarma et al. (2020) found that increased salinity exposure decreased the growth and survival rates in *Labeo rohita*, with maximum mortality occurring at 4.5 ppt. In *Pangasianodon hypophthalmus*, Hossain et al. (2021) observed that the LC50 salinity concentration for embryos was 11.24ppt, while for larvae it was 10.63ppt. These various studies provided valuable insights into the salinity tolerances of various fish species and contributed to our understanding of the complex relationship between fish and salinity, with the importance of considering salinity as a crucial factor in habitat management and conservation efforts for freshwater and brackish water species.

Different salinity regimes have been shown to affect gene expression levels, leading to alterations in the physiology of organisms. These changes can be elucidated through gene expression studies. In the present investigation, the expression of IGF mRNA was examined in the gills of pearl spot fish at various salinity levels. It was found that at 15ppt, the expression of IGF mRNA was upregulated compared to the control, but as salinity increased further, the expression began to decline. The peak down-regulation was observed at 45ppt. These findings align with previous research conducted by Beckman (2011) on striped sea bass, a euryhaline teleost. Beckman identified a decrease in liver IGF-I mRNA levels following the transfer of fish from freshwater to seawater, suggesting that a decline in hepatic IGF-I production may be responsible for the reduced plasma levels. In another study involving Mozambique tilapia, a euryhaline fish native to estuary environments, Seale et al. (2020) provided insights into the sex-specific modulation of growth hormone (GH/IGF) expression by environmental variables. They reported that compared to freshwater conditions, tilapia exhibited enhanced growth and greater GH expression in tidal regimes with varying salinity. Furthermore, Breves et al. (2010) conducted a separate investigation and discovered that prolonged exposure to salinity resulted in a decrease in IGF expression in Mozambique tilapia, while GH hormone expression showed a slight increase during the transfer to seawater. Breves et al. (2011) noted increased PRL and GH receptors in osmoregulatory tissues in FW-acclimated *Oreochromis mossambicus*. The mRNA expression of IGF-1 in pearl spot gill indicated that the fish perform well and show good growth at 15ppt water which supports the fact that pearl spot is a brackish water euryhaline fish which can withstand a wide range of salinity (Arun kumar et al., 2020). Fish require energy to maintain their metabolic functions, including osmoregulation and oxygen consumption. The energy required for these processes increases as the salinity of the water increases. However, fish reared in isosmotic or less saline water need to expend less energy on these metabolic functions, resulting in a conservation of energy that can be utilized for other activities, such as growth. This is one of the explanations for the increased IGF expression observed in fish reared in less saline water. By diverting energy away from osmoregulation and towards growth, fish can enhance their chances of survival in suboptimal environments. This phenomenon has been observed in numerous studies, including those examining the effects of salinity on IGF expression in pearl spot fish and Mozambique tilapia. The understanding of how energy is utilized in fish can provide insight into the physiological mechanisms underlying their responses to environmental changes and can be valuable in developing strategies to improve fish farming practices and fisheries management.

Heat shock protein (HSP) is widely recognized as a gene that responds to stress, and a lower level of HSP expression often indicates a relatively stress-free condition (Sanders, 1993). The findings of the current study on pearl spot fish support this theory, as they revealed that IGF-1 mRNA expression reached its highest level while HSP70 mRNA expression was at its lowest point in salinity of 15ppt. Moreover, there appeared to be an inverse correlation between HSP70 and IGF-1 gene expression, with HSP70 mRNA levels increasing as salinity levels increased. The peak expression of HSP70 mRNA was finally observed in the hypersaline condition of 45ppt, exhibiting a remarkable fold change value of 7.3. Comparable results were observed in black sea bream by Deane et al. (2002), who reported an inverse association of expression between IGF-1 and hsp90 genes under isotonic environmental conditions (12ppt). Similar findings were also reported by McCormick (2001), where the roles of GH/IGF, prolactin, and cortisol in osmoregulation were investigated. GH/IGF was found to promote acclimation to seawater, prolactin to freshwater acclimation, and cortisol to collaborate with both hormones, providing a dual osmoregulatory function. Additionally, in carp, exposure to pond water containing 1% wt/vol sodium chloride led to a significant increase in hepatic HSP70 levels (De Wachter et al., 1998). These collective findings demonstrate the dynamic relationship between salinity, stress response genes such as HSP70, and other regulatory factors involved in osmoregulation and growth. Understanding the intricate interactions between stress-indicating genes, growth factors, and hormones in response to salinity fluctuations contributes to our knowledge of fish adaptation mechanisms.

The migration of euryhaline teleost species across various salinity gradients relies on the activity of Na+/K+-ATPase (NKA), as demonstrated in this study. The research highlights the remarkable adaptability of pearl spot to a wide range of salinities. Specifically, the study observed a significant upregulation of NKA gene expression in the gills at 15ppt, distinguishing it from the expressions at 25, 35, and 45ppt. Previous investigations by Tipsmark et al. (2011), revealed the existence of two NaKATPase-subunit isoforms (NKA-1a and NKA-1b) in *Mozambique tilapia* and observed that during acclimatization from freshwater to seawater, NKA-1a expression decreased considerably within 24 hours, while NKA-1b expression increased. Likewise, in the case of transferring seawater-acclimated fish back to freshwater, there was an initial increase in NKA-1a expression within two days, followed by a subsequent decrease in NKA-1b expression after 14 days. The current study found that NKA expression in pearl spot is similar to NKA-1a expression, and a homology search using NCBI Blast confirmed that the NKA gene sequence in the present study is Type 1a. Another study by Chandrasekar et al. (2014) reported two isoforms of NKA with distinct expression patterns during water acclimatization to different salinities in pearl spot. One isoform exhibited increased abundance in response to seawater acclimation, indicating a role in ion secretion similar to NKA a1b, while the expression of the other isoform increased during both freshwater and seawater acclimation, suggesting isoform switching during salinity acclimation. Additionally, Liang et al. (2017) investigated the effects of salinity on the expression of NKA genes in the gill tissue of *Anguilla marmorata*, a catadromous euryhaline species. They observed a gradual increase in NKA expression in brackish water (BW) and seawater (SW) compared to freshwater (FW), reaching peak levels at 12 and 24 hours post-transfer with 3.6-fold and 8.4-fold increases, respectively. Maintaining intracellular ion concentrations within specific physiological ranges is vital for an organism’s normal biological function, which consequently leads to a reduction in NKA expression (Jia and Liu, 2018). Overall, these findings shed light on the complex interplay between NKA expression, salinity adaptation, and the temporal aspects of salinity exposure, furthering our understanding of the mechanisms underlying osmoregulation in aquatic organisms.

Khodabandeh et al. (2008) reported on the acclimation of *Liza aurtata* fry to different salinities, including freshwater (FW) and 12, 36, and 46ppt. The intensity of NKA showed no significant change as salinity increased from 36 to 46ppt, but it was notably higher (p>0.05) in FW compared to salinity of 12ppt. Gill NKA activity exhibited a significantly higher level (p>0.05) in fish acclimated to 36ppt and 46ppt salinity (3.3- and 5.1-fold) than in fish acclimated to 12ppt salinity. Other studies have explored the role of regulatory subunits, such as FXYD protein and claudins, in osmotic homeostasis and permeability changes associated with salinity acclimation. Further investigations could establish interactions between FXYD and NKA, shedding light on the broader molecular mechanisms involved in salinity adaptation in euryhaline fish (Wang et al., 2008; Tipsmarks, 2008; Tipsmarks et al., 2008; Tipsmarks et al., 2016). Overall, these studies contribute to our understanding of the complex processes underlying osmoregulation and highlight the intricate molecular responses of aquatic organisms to salinity challenges.

Oxidative stress, indicated by an excess of reactive oxygen species (ROS), is a critical factor affecting organismal health. The study investigated the expression patterns of Superoxide dismutases (SODs) and Catalase Activity (CAT) in the liver of pearl spot at different salinities. The findings revealed that both SOD and CAT exhibited an increase in expression at 25 and 35ppt salinities, followed by a decline at higher salinities. These results align with similar studies conducted in other fish species, such as silver pomfret and marbled eel, which demonstrated an augmentation of antioxidant activity in response to increasing salinity to counteract ROS-induced damage (Yin et al., 2011; Wang et al., 2016). Additionally, studies in *Fundulus heteroclitus* and *Labeo rohita* further support the correlation between salinity and antioxidant defences, indicating an increase in total oxidative scavenging capacity (TOSC) and SOD/CAT activities with rising salinity levels. However, it is noteworthy that while antioxidant defences tended to increase, Na+/K+-ATPase (NKA) activity decreased as salinity increased, consistent with the current study’s findings. These findings emphasize the intricate interplay between salinity, oxidative stress, and physiological responses, highlighting the importance of antioxidant defences as early detection biomarkers for stress-induced adverse effects in organisms exposed to changing environmental conditions.

Gene expression changes play a crucial role in mediating the physiological acclimations of euryhaline fish to osmotic stress. The current study focused on investigating the mRNA expression levels of OSTF1, an inducible transcription factor associated with osmotic stress response, in the gills of pearl spot at different salinities. The results demonstrated that at 25ppt salinity, there was a significant upregulation of OSTF1 mRNA expression, indicating its responsiveness to osmotic stress conditions. However, as the duration of the stress increased, the expression level of OSTF1 decreased at 35 and 45ppt salinities. These findings are consistent with previous studies conducted on tilapia and milkfish (Fiol et al., 2006; Tse et al., 2014), which also observed rapid and transient induction of OSTF1 during osmotic stress. The collective evidence supports the role of OSTF1 as an important component in osmosensory signal transduction and transcriptional regulation for osmotic adaptation in euryhaline teleosts. The quick responsiveness of OSTF1 to osmotic stress, as observed in pearl spot and other fish species, highlights its critical involvement in fish osmoregulation. Additionally, the link between OSTF1 and cortisol, as established by Lin et al. (2022) further emphasizes the complexity of the osmotic stress response, indicating that cortisol can enhance the expression levels of both OSTF1a and OSTF1b. These findings shed light on the regulatory mechanisms underlying osmotic stress adaptation in fish and emphasize the importance of OSTF1 as an early and primary responsive gene during hypertonic stress. Understanding the transcriptional regulation mediated by OSTF1 and its interaction with other factors, such as cortisol, provides valuable insights into the molecular pathways involved in fish osmoregulation.

The comprehensive analysis of gene expression values in response to varying environmental salinity provides valuable insights into the adaptive mechanisms of fish species, specifically pearl spot, to osmotic stress. The findings highlight the dynamic nature of gene expression changes and their crucial role in mediating physiological acclimations and osmoregulation.

### 4.1. Histological Changes in Eutroplus suratensis at Varying Salinity

The histological examination of gill tissues provides valuable insights into the adaptive responses of pearl spot to varying salinities, shedding light on the morphological changes that enable their osmoregulatory capabilities. In chloride cells, higher salinity levels can lead to significant physiological dysfunction and respiratory damage. The present study investigates the morphological alterations observed in pearl spot gills during adaptation to a wide range of salinities, utilizing histological analysis. At higher salinity levels (25, 35, and 45ppt), severe physiological damages were observed, including lamellar curling, enlargement of lamellae and chloride cells, necrosis, epithelial haemorrhage, blood congestion, and pronounced swelling in the lamellar region, known as severe edema. In contrast, salinities ranging from 0 to 15ppt did not exhibit any significant effects. These findings highlight the sensitivity of chloride cells to higher salinity and the resulting detrimental consequences on the overall physiological integrity of pearl spot. Similar histological changes were observed by Kumar et al. (2020) in pearl spot acclimated to freshwater (FW), brackish water (BW), and seawater (SW). Compared to SW-acclimated fish, those acclimated to FW and BW displayed fewer chloride cells in the inter-lamellar region. Increased salinity in SW induced physiological changes, including an expansion of chloride cells and the lamellar region. The severe respiratory challenges experienced in SW conditions resulted in lamellar hyperplasia, necrosis, lamellar fusion, lamellar curling, and epithelial rupture with haemorrhages, all indicative of osmotic stress-induced damage. The osmoregulatory capacity of teleosts varies depending on the integrated functions of organs such as the kidney, digestive tract, and gills (Cioni et al., 1991). Studies have demonstrated that the number and size of chloride cells in the gills of euryhaline teleosts, including *Oreochromis sp*., increase upon transfer from FW to SW (Avella et al., 1993). In pearl spot, an increase in salinity from 15 to 35ppt (from brackish water to seawater conditions) led to a further decrease in the number of mucus cells, consistent with observations made by Virabadrachari (1961) in *Etroplus maculatus*. Lamellar hyperplasia, resulting from the proliferation of lamellar cells, occurs during hyperplastic conditions and contributes to a reduction in inter-lamellar space, potentially leading to lamellar fusion (Fracário et al., 2003). Respiratory system disruption in fish due to increasing salinity concentrations has been identified as one of the contributing factors to lamellar hyperplasia (Ried et al., 2006). Shahriari Moghadam et al. (2013) investigated the primary target organ for acute salinity effects in Liza Aurata and reported that it affects the type of chloride cells and gill structure.

The collective findings from these histological studies emphasize the intricate morphological responses of pearl spot gills to varying salinities and highlight the challenges imposed by osmotic stress. Understanding these adaptive changes in gill tissues contributes to our knowledge of osmoregulatory mechanisms in fish and provides valuable insights for assessing the effects of environmental stressors on fish populations. Further research in this area is warranted to deepen our understanding of the underlying physiological and cellular mechanisms involved in osmoregulation.

## 5. Conclusion

This study aimed to investigate the differential expression of growth, stress, and ion regulatory genes in pearl spot under different salinity conditions. Through assessing the lethal concentration of salinity through LC50 the pearl spot fish were acclimatized to varying salinity concentrations. qRT-PCR was employed to assess the differential expression of selected genes associated with ion regulation (Na+/K+-ATPase), growth (IGF-1), and stress response (HSP, catalase, and SOD). The results indicated that the growth-related gene IGF-1 was significantly upregulated at 15ppt and downregulated at 45ppt, suggesting that lower salinities promote pearl spot growth. The expression of IGF-1 was found to be linked to the HSP70 gene, which exhibited higher expression levels at higher salinities, indicating oxidative stress. The OSTF-1 gene was expressed in response to osmotic stress, with acute and rapid upregulation at 15ppt, peaking at 25ppt and 35ppt, and downregulation at 45ppt. NKA activity in the gills increased at 15ppt and downregulated at 35ppt, suggesting that 15ppt was the tolerance limit for ion balance regulation. Analysis of SOD and CAT mRNA levels in the liver showed upregulation at 25ppt, indicating oxidative stress beyond the fish’s tolerances. Histology examinations revealed significant physiological changes in pearl spot gills at salinities of 25ppt, 35ppt, and 45ppt. These changes included edema in the lamellar region, blood vessel congestion, lamellar curling, lamellar fusion, necrosis, and enlargement of chloride cells. These findings contribute to our understanding of the molecular basis of osmotic stress adaptation in fish species and provide potential biomarkers for monitoring environmental stressors. The knowledge gained from these studies can help inform conservation and management strategies for euryhaline fish species, ensuring their resilience in the face of changing salinity environments. Further research in this field holds promise for unraveling additional molecular mechanisms and refining our understanding of osmoregulation in fish.

## Acknowledgement

This work was carried out under the project (NICRA) National Initiative on Climate Resilient Agriculture receiving funding support from the (ICAR) Indian Council of Agricultural Research. The author would like to thank Tamil Nadu Dr. J. Jayalalithaa Fisheries University (TNJFU) and **T**NJFU **-** Institute of Fisheries Post Graduate Studies, Vaniyanchavadi, Chennai for providing facilities for carrying out this work.

